# A Rapid Pathogenicity Assay of *R. solanacearum* in 10-12 Days Old Eggplant (*Solanum melongena*) Seedlings by Root Inoculation Approach

**DOI:** 10.1101/2020.04.08.033050

**Authors:** Niraj Singh

## Abstract

This study has reported standardization of pathogenicity assay of *Ralstonia solancearum* in eggplant seedlings by root inoculation approach. Though the approach is similar to my earlier published work that done with tomato seedlings, in case of eggplant, duration of the seed germination, age of the seedlings, symptom timing and progression rate of the disease are different. In an introductory observation, *R. solanacearum* F1C1 inoculation was performed in to 10-12 days old eggplant seedlings resulted in lethal infection in eggplant seedlings and subsequently most of these infected seedlings got wilted within 10 days of post-inoculation. Colonization studies of *R. solanacearum* in wilted as well as infected eggplant seedlings were confirmed by GUS staining as well as fluorescence microscopy. In addition, we also observed that wrapping of the eggplant seedlings root by a thin layer of cotton soon after the pathogen inoculation in the root, enhanced the disease progression and wilting of the inoculated seedlings. The standardized root inoculation protocol in eggplant seedlings was found to be efficient to distinguish mutant strains of different virulence genes such as *hrpB, phcA* and *pilT* from wild-type *R. solanacearum*. Due to its reproducibility and consistency in different eggplant cultivars, the standardized protocol described here is practically observed as an useful and rapid approach to investigate *R. solanacearum* pathogenicity and disease progression at the early seedling stage of eggplant.

## 2. Introduction

*R. solanacearum* is a soil-born bacterium, the causal agent of bacterial wilt disease in many plant species including eggplant and other vegetable crops (Wicker et al., 2007; Genin, 2010). *R. solanacearum* infection and wilting is not only localized to its specific host, but also it spreads from one host to the other through soil, contaminated water and weeds (Genin, 2010). In spite of broad host range of this pathogen, only few host plants such as tomato, potato and *Arabidopsis thaliana* have been used mainly for studying pathogenicity. Similar to the tomato plants, bacterial wilt is also a very common disease in eggplants in tropical as well as in subtropical regions (Gaitonde and Ramesh; Smith, 1896; Shekhawat et al., 1978; Ramesh et al., 2009; Salgon et al., 2017). Eggplants one of the most common vegetables, consumed in various ways by a large population across the world and it is a rich source of protective nutrients and fibers (Hedges and Lister, 2007). Eggplant belongs to solanaceae family, it is an agriculturally important crop plant for pathogenicity and bacterial wilt related study. Till now, determination of *R. solanacearum* pathogenicity factors in case of eggplant has not been addressed appropriately in comparison to tomato or some other solanaceae crop. Studies on *R. solanacearum* pathogenicity in eggplants have mainly been to screen resistant cultivars against the bacterial wilt disease and to investigate resistant mechanisms of eggplant against the wilt pathogen (Artal et al., 2013; Gopalakrishnan et al., 2014).

In addition to that, most of solanaceae family crop plant species such as tomato, potato, and pepper *etc*. have originated from the new world, while eggplant and its associated wild relatives belongs to the old world. It is being assumed that eggplant was domesticated in Southeast Asia region (Daunay, 2008; Syfert et al., 2016). While on the genetic level, both tomato and eggplant are autogamous diploid plant species with twelve chromosomes and nearly equal genome size (Arumuganathan and Earle, 1991), therefore, deep investigation and study on co-evolution between eggplant and its associated old world pathogen relationship may point out why some specific resistance genes do not exist in the new world solanaceous crops plant (Hirakawa et al., 2014; Nahar et al., 2014). Along with its agronomical importance, eggplant genomics are also less studied and documented in comparison to other major solanaceous crops like tomato and potato. A study was also carried out in eggplant under controlled environment against a core collection of twelve strains, representative of all the exiting phylotypes of *R. solanacearum* (Lebeau et al., 2011). The result of that study indicated that eggplant is a potential host for most of the existing phylotypes of *R. solanacearum*. Thus, pathogenicity study in eggplant as well as comparative virulence and disease progression study between other common hosts like tomato, eggplant might be helpful for in-depth understanding on molecular mechanism of plant-microbe interaction during infection and wilting. This study can also suitably project eggplant seedling as a suitable model for mapping and investigating resistance factors to pathogens as well as other determinates. On the basis of severity of wilt disease and economic importance of this crop, the present investigation of *R. solanacearum* pathogenicity behavior and disease progression study in eggplant seedlings will be very helpful to generate necessary information for the understanding of eggplant and *R. solanacearum* interaction during infection and wilting. Over and above, this study will be also very significant for the economic as well as scientific interest for suitable management measures to minimize the substantial crop loss.

## 3. Materials and Methods

### 3.1 Bacterial strains, growth media and culture conditions

*R. solanacearum* as well as other bacterial strains recruited in this work were the same strains which have been listed in the recently published tomato seedlings work (Singh et al., 2018). Growth medium and antibiotics were also used as same as mentioned in the tomato seedlings work. *R. solanacearum* F1C1 (Kumar *et al*., 2013) and its derivative mutant strains as well as *Pseudomonas putida* were grown in BG medium (Bacto glucose agar medium) (Boucher *et al*., 1985) supplemented with glucose (0.5%, w/v). Incubation temperature for *R. solanacearum* strains and *Pseudomonas putida* was 28°C while *Bacillus subtilis* as well as *Escherichia coli* strains were grown in LB medium (Bertani, 1952) at 37°C in the same way as discussed in case of tomato seedlings. Agar (1.5%, w/v) was added in case of solid medium if and when necessary.

### 3.2 Germination of eggplant seedlings for inoculation

Seeds of different eggplant cultivars employed during pathogenicity assay standardization were pre-soaked in sterile distilled water for 5-6 days. Further, this was followed by spreading of the soaked seeds on sterile and wet tissue paper bed in a plastic tray. After that, plastic tray was covered with transparent polyethylene sheet and allowed for germination in a growth chamber (Orbitek, Scigenics, India) maintained at 28°C, 75% relative humidity (RH) and 12 h photoperiod respectively. It usually took 2-3 days for sprouting of seeds. Just after sprouting, polyethylene sheet cover was removed from the tray, sterile distilled water was sprinkled at regular time interval till 7-8 days to sustain the germination process and growth of seedling in a better way. Here the age of the eggplants seedlings have been defined from the day of spreading the soaked seeds for germination on the wet tissue paper bed in the plastic tray.

### 3.3 Preparation of bacterial inoculums

*R. solanacearum* F1C1 as well as other bacterial strain inoculums were prepared by following exactly the same procedure used in case of tomato seedlings inoculation as mentioned in case of tomato seedling(Singh et al., 2018).

### 3.4 Root inoculation of *R. solanacearum* in eggplant seedlings

In case of eggplant, around 15-20 ml of *R. solanacearum* F1C1 inoculum (∼ 10^9^ CFUml^-1^) was taken in a sterile container similar to tomato seedling inoculation as mentioned in case of tomato (Singh et al., 2018). After that, from the germinated seedling bed, 10-12days old eggplant seedlings were picked one at a time. Root of each seedling was then carefully dipped (up to the root-shoot junction) into the bacterial inoculum for 1-2 seconds followed by transfer of the seedling to an empty and sterile microfuge tube of 1.5 or 2.0 ml (Figure: 1A & 1B). All the eggplant seedlings were inoculated by the same procedure as mentioned in earlier infection assay in tomato seedlings.

**Figure 1A:**
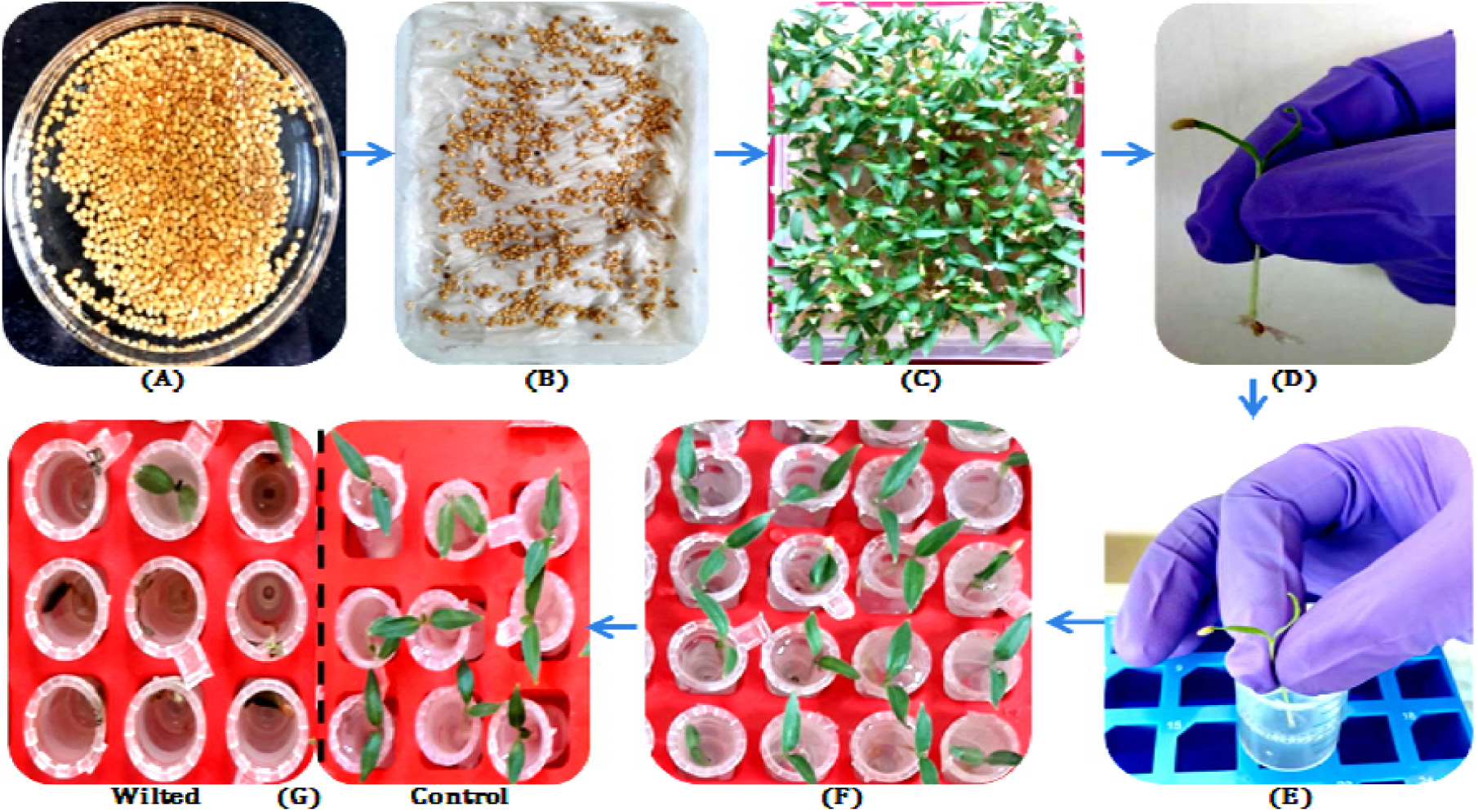
Pictorial presentation of root inoculation method in eggplant seedlings. **(A)** Soaking of eggplants seeds in sterile distilled water, **(B)** Germination of eggplant seeds on cotton bed, **(C)** Germinated seedlings in plate after 10-12 days, **(D)** A 10-12 days old eggplant seedling, **(E)** Root inoculation of eggplant seedling in pathogen inoculum, **(F)** Transfer of each inoculated seedling to sterile empty microfuge (1.5–2.0 ml) tubes and after five minutes exposure to air, sterile water is added in the microfuge, **(G)** After 10 days, 80%-90% of infected tomato seedlings were wilted/died (left side) in comparison to water as control (right side).

**Figure 1B:**
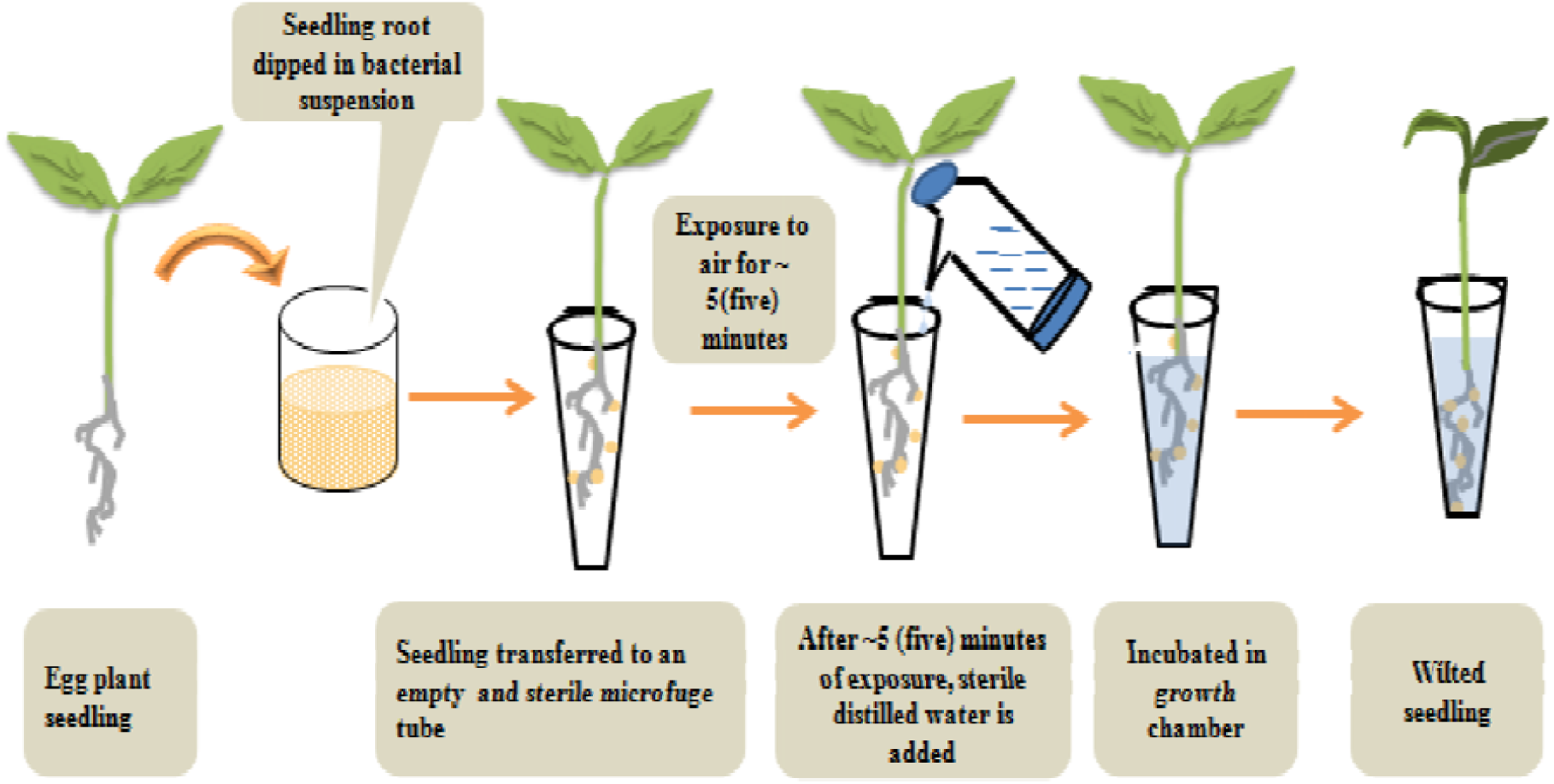
Schematic representation of the crucial steps of root inoculation method in eggplant seedlings. A schematic representation emphasizing the exposure to air after pathogen inoculation and various steps during the pathogenicity assay of *R. solanacearum* in 10-12 days old eggplant seedlings by root inoculation method

In all these experiments, a set of forty eggplant seedlings was considered for each pathogenicity test per inoculum used. In control set also, 40 (forty) seedlings were mock-inoculated with sterile distilled water. All the inoculated eggplant seedlings along with control were transferred to a growth chamber maintained at 28°C, 75% RH, photoperiod of 12 h. The inoculated seedlings were analyzed for disease progression and wilting from next day onwards (inoculation) till 10^th^ day of post-inoculation and findings were recorded accordingly.

During this work, seedlings of four different commercially available eggplants cultivars such as Akshay (Bharat nursery Pvt. Ltd.), Lalita (Bharat nursery Pvt. Ltd), Debgiri (Debgiri Agro Products Pvt. Ltd.) and 749 (Pahuja Seed Pvt. Ltd.) were tested for *R. solanacearum* pathogenicity and disease progression by the root inoculation method as described above.

Root inoculation of *hrpB, phcA, pilT* and *rpoN*_*2*_ mutant strains of *R. solanacearum* as well as non-pathogenic bacterial strains such as *B. subtilis, P. putida*, and *E. coli* in eggplant seedlings were carried out in the same way as mentioned in case of tomato seedlings (Singh et al., 2018).

### 3.5 Colonization study of *R. solanacearum* in eggplant seedlings

Grown culture of TRS1002 (GUS marked), TRS1016 (mCherry marked)(Monteiro et al., 2012; Capela et al., 2017) as well as TRS1017 (GFP marked) strain of *R. solanacearum* were pelleted down by centrifugation and 10^9^CFU/ml inoculum was prepared by the same procedure described for *R. solanacearum* in tomato seedlings work (Singh et al., 2018). Thereafter, bacterial inoculums were used for infection by root inoculation approach in 10-12 days old eggplant seedlings in the same way as stated above. After third day of post inoculation, eggplant seedlings were employed for colonization study followed by the same procedure as mentioned in the case of tomato seedlings.

### 3.6 Study of cotton fiber wrapping on the inoculated root of eggplant seedling and its impact on wilting

During the standardization process of *R. solanacearum* pathogenicity assay in early seedling stage of eggplant, this study has gone through many trial and error among the different steps of pathogenicity standardization process. During standardization it was observed that a thin layer of cotton fiber if wrapped on the root of eggplant seedling just after the pathogen inoculation, it used to enhance the wilting and disease progression as compared to those set of seedlings without the wrapping of cotton fiber (Figure 2). During, pathogen inoculation process, we have performed in the same way as motioned above, after that a thin layer of sterile cotton fiber was wrapped around the seedling root up to root shoot junction (Figure 3). After that, infected seedlings were transferred to sterile microfuge tube and allowed for ∼5 (five) minutes of air exposure. Thereafter, water is added to the microfuge in the same way as described in the above sections as well as in the previous chapter.

**Figure 2:**
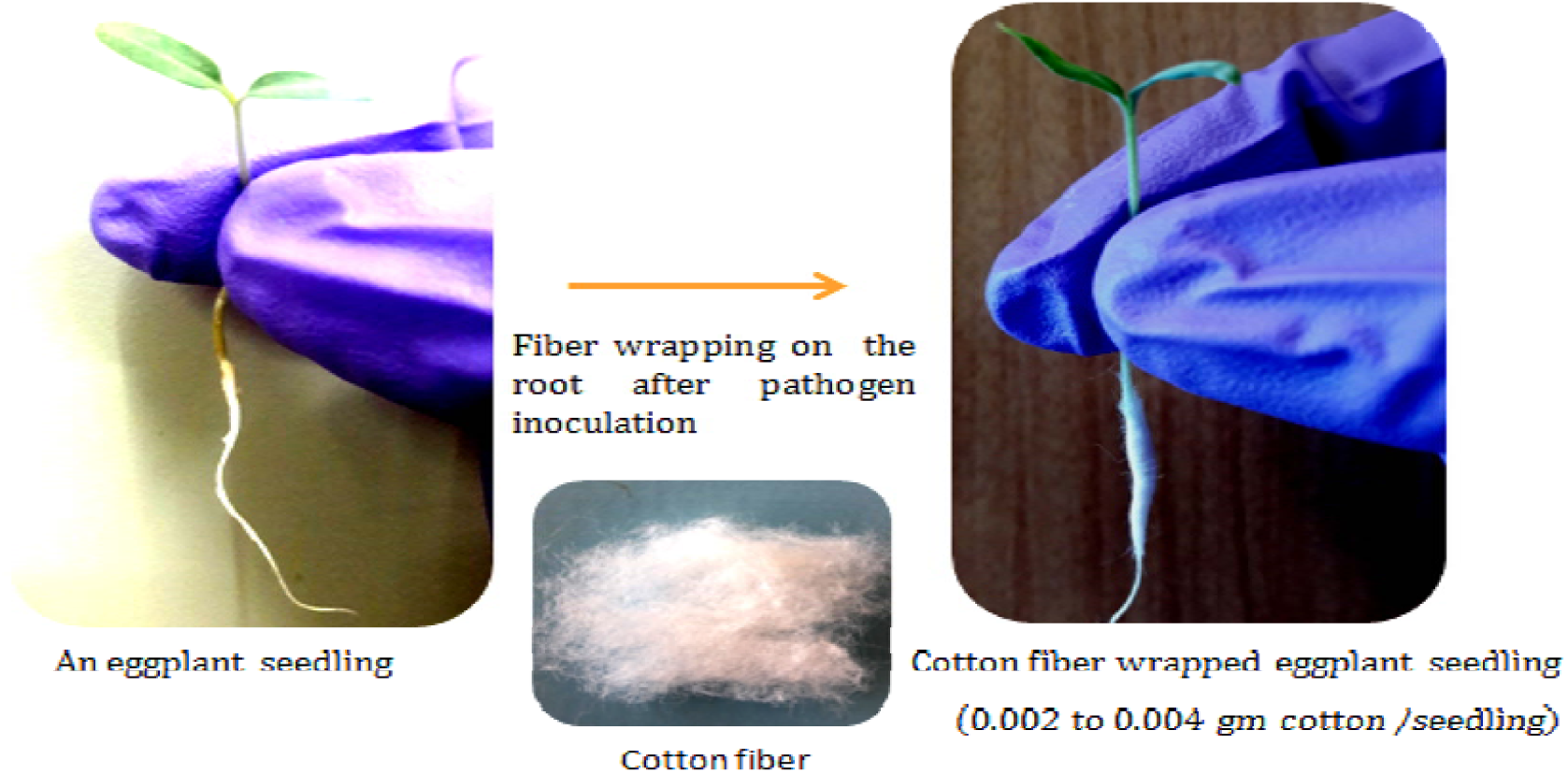
Pictorial presentation of the steps of cotton wrapping on the root of eggplant seedlings. This figure is showing the 10-12 days old eggplant seedling before and after cotton fiber wrapping process on root part of eggplant seedling just after the pathogen inoculation process.

**Figure 3:**
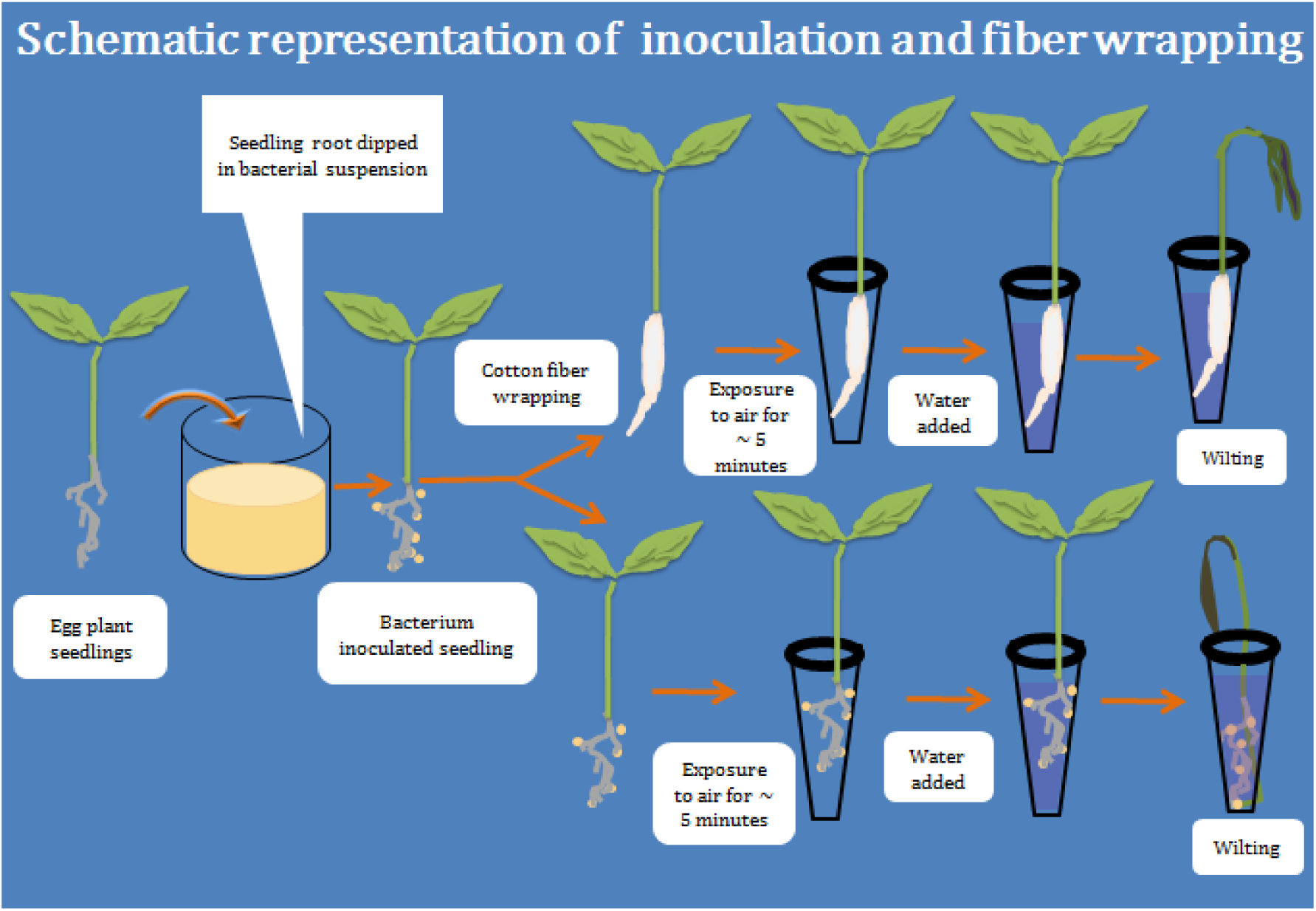
Schematic representation of pathogen inoculation and cotton fiber wrapping on the root of eggplant seedlings. A self-explanatory sketch describing how sterile cotton was wrapped around the root of eggplant seedling just after pathogen inoculation step. The whole process was used to understand the impact of cotton fiber on *R. solanacearum* wilting in the eggplant seedlings by comparing pathogenicity between cotton wrapped and unwrapped set of infected eggplant seedlings.

## 4 Results

### 4.1 *R. solanacearum* pathogenicity in eggplant seedlings by a root inoculation method

In the earlier published work, development of pathogenicity assay and wilt disease progression in 6-7 days old tomato seedlings were studied and susceptibility of early stages of tomato seedlings toward *R. solanacearum* wilting was also validated. In addition to that, this early seedling stage pathogenicity study approach was found to be very much consistent and proficient. In wake of testing reproducibility of this newly standardized pathogenicity approach in eggplant seedling as another most affected host, we have carried out this work. In the inoculation process, we have followed the same steps as mentioned in tomato seedlings. For pathogenicity standardization, 10-12days old eggplant seedlings were inoculated with *R. solanacearum* suspension, thereafter inoculated seedlings sets along with controls were transferred to growth chamber for the observation of disease progression. On the 10^th^day post-inoculation, about 80-90% of the inoculated eggplant seedlings were found to be dead due to disease and wilting (Figure 4).

**Figure 4:**
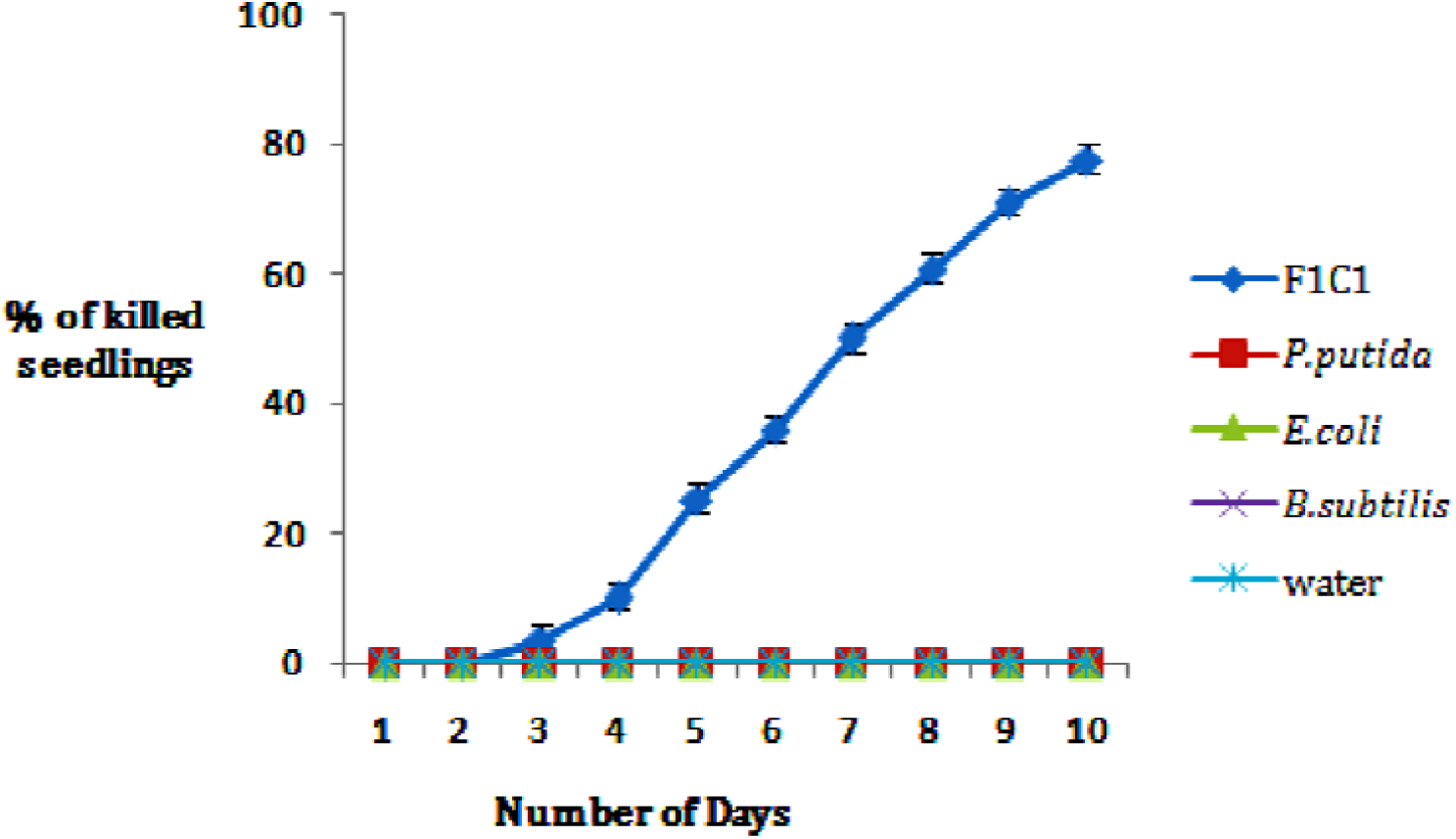
Pathogenicity assay of F1C1 and other bacterial strains in eggplant seedlings. This graph is showing the wilting or death occurred only in F1C1 infected eggplant seedlings. In case of *P. putida, B. subtilis* and *E. coli*, no killing was observed which was similar to water as control.

For further confirmation of the wilting of inoculated eggplant seedlings by root inoculation approach specific to *R. solanacearum*, eggplant seedlings were also inoculated with some well-known non-pathogenic bacteria such as *P. putida, B. subtilis*, and *E. coli* along with sets of *R. solanacearum* infection (Figure 4). After 10^th^day of post-inoculation, it is observed that none of the eggplant seedlings exhibited disease symptom inoculated with these nonpathogenic bacteria. This result of infection clearly indicated that wilting and death of the eggplant seedlings occurred only due to inoculation with *R. solanacearum* F1C1. Further, to confirm reproducibility of this standardized root inoculation approach, it was again tested on three different eggplant cultivars such as Lalita, Debgiri and 749. Magnitude of *R. solanacearum* F1C1 pathogenicity could be observed as similar kind in all three eggplant cultivars (Figure 5 & 6).

**Figure 5:**
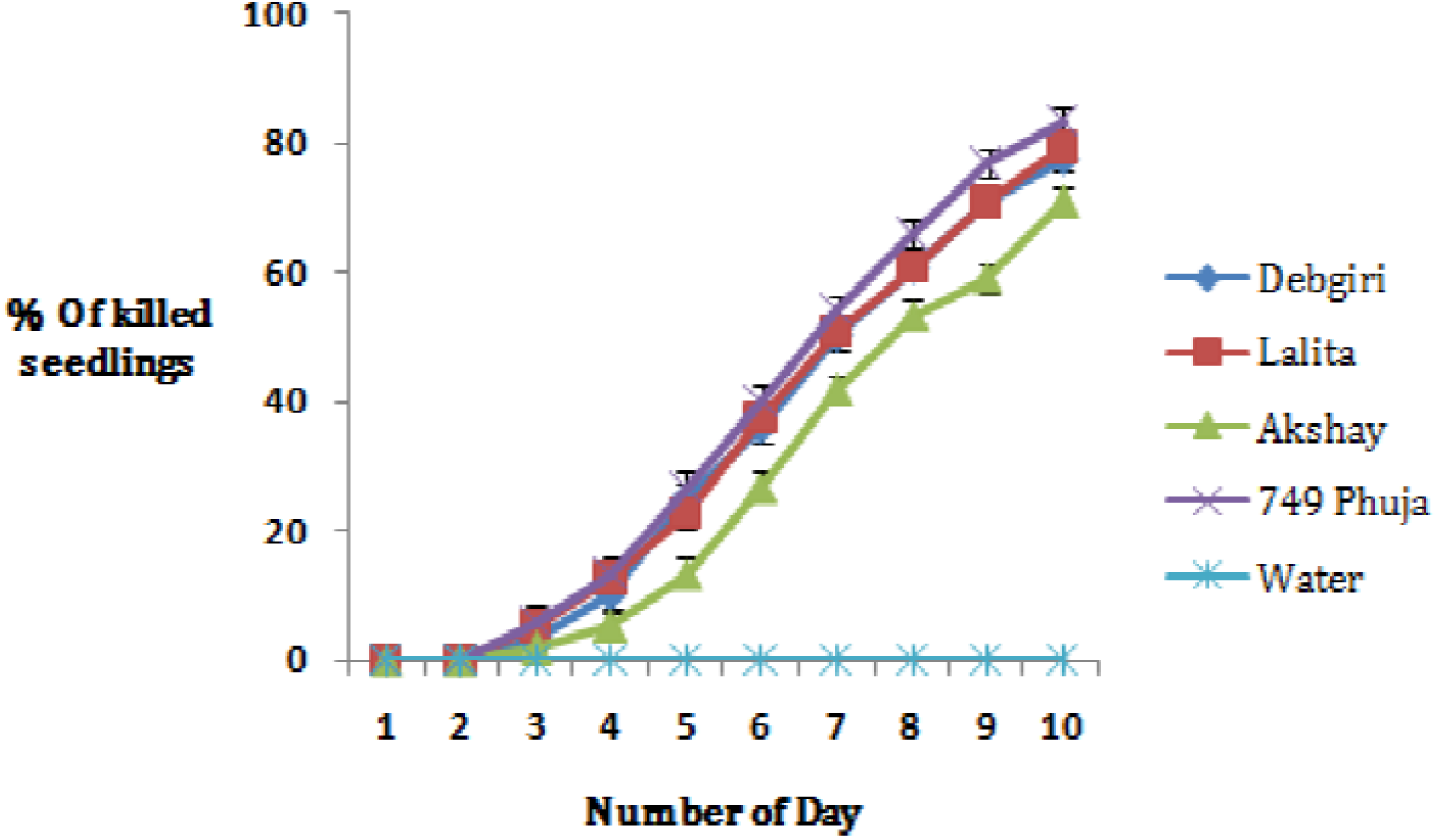
Pathogenicity assay in different eggplant cultivars. This graphis showing the pathogenicity assay of *R. solanacearum*F1C1 in four different eggplant cultivars by root inoculation method. It can be clearly observed that this *R. solanacearum* F1C1 pathogenicity by this inoculation method was nearly equal to virulence and disease progression in all the four different eggplant cultivars.

**Figure 3.6:**
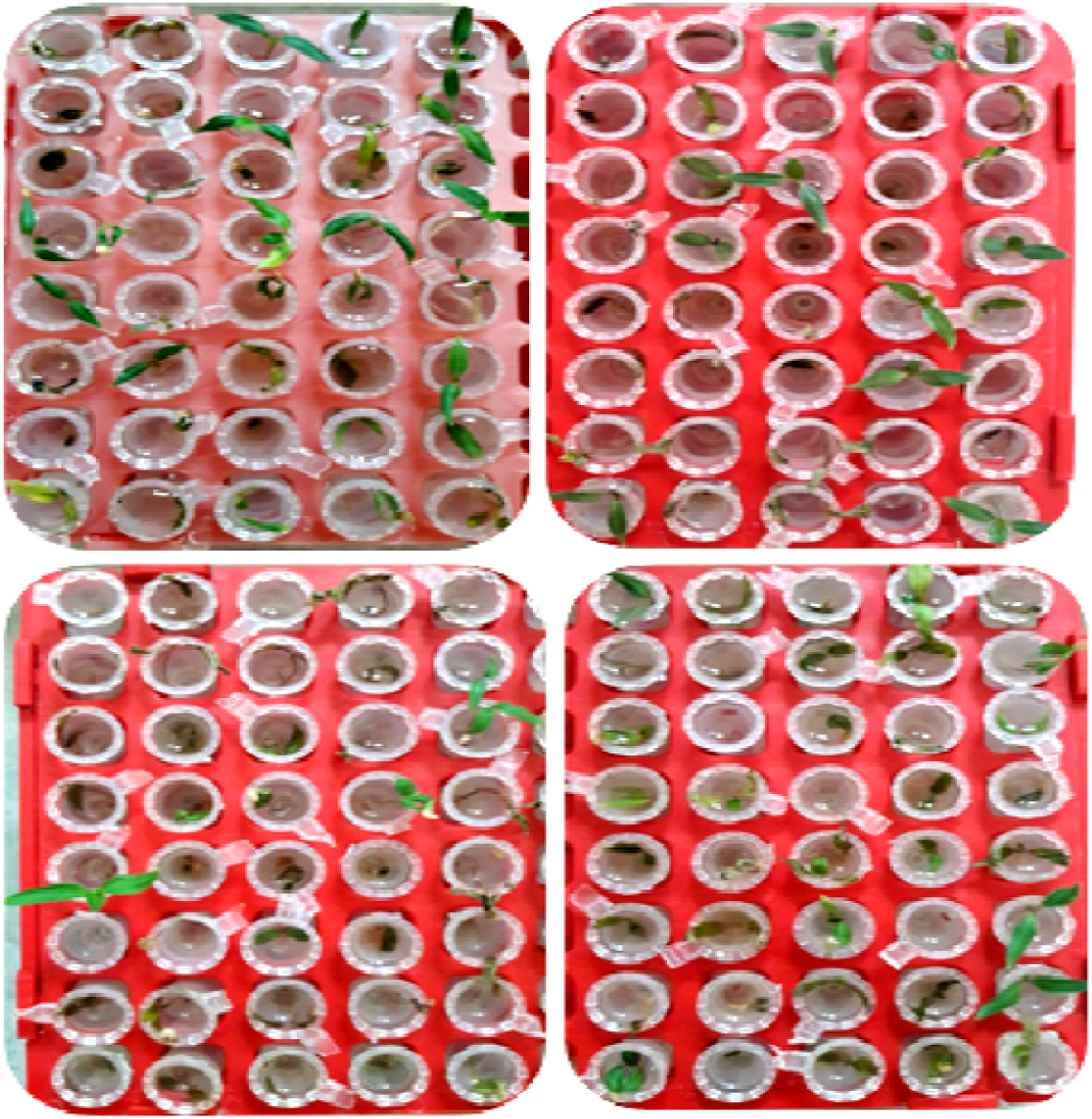
Pictorial presentation of pathogenicity assay in four different eggplant cultivars by root inoculation. These pictures are showing the pathogenicity assay of *R. solanacearum* F1C1 in four different eggplant cultivars by root inoculation method. It can be clearly observed that this *R. solanacearum* F1C1 pathogenicity by root inoculation method was highly reproducible in all the four different eggplant cultivars. in left side upper and lower picture indicating Akshay and 749 Pahuja cultivar while in right side upper and lower picture indicating Debgiri and Lalita. Wilting and disease progression was found to be consistent in respective cultivar.

### 4.2 Colonization study

Colonization study in eggplant seedling was carried out by using *R. solanacearum* F1C1 derived strains such as TRS1002 (GUS marked), TRS1016 (mCherry marked) as well as TRS1017 (GFP marked). Bacterial colonization in the infected seedlings was observed from root to the shoot regions, which were confirmed by GUS staining as well as by red fluorescence (Figure .7). This result of colonization investigation suggested the growth and colonization of the pathogen from root to the shoot regions during the infection process and disease progression.

**Figure 7:**
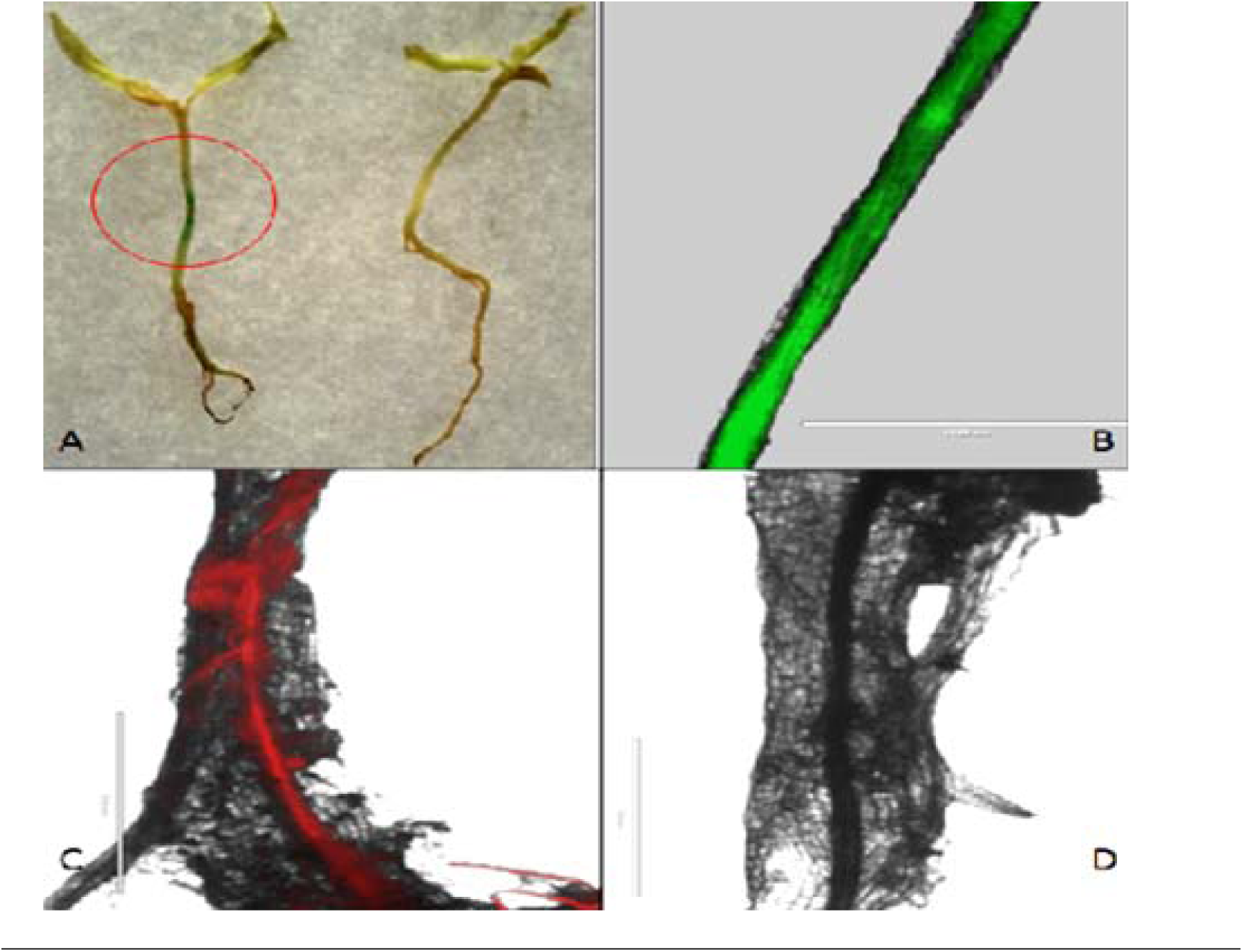
Pathogen colonization study with the help of GUS staining and fluorescence microscopy in eggplant seedlings. (A) is showing GUS staining (on the stem) for colonization confirmation (red circle) while right side seedling iscontrol showing no blue color, (B) is showing green fluorescence in root region which is confirming the colonization of GFP tag *R. solanacearum* strain, (C) is showing the red fluorescence in root region which is confirming the presence and colonization of m-cherry tagged *R. solanacearum* inside tomato seedling while (d) is showing the root region of control seedling with the absence of any florescence.

### 4.3 The root inoculation method can be used to study *R. solanacearum* and its mutant derivatives virulence functions

To further validate and find out the usefulness of this standardized root inoculation protocol to study the pathogenicity functions of *R. solanacearum* eggplant seedlings, seedlings root were inoculated with different *R. solanacearum* mutant strains such as *hrpB* (TRS1012), *phcA* (TRS1013), *pilT* (TRS1014) and *rpoN2* (TRS1015) in the similar way as mentioned in the previous Seedling published work. As assumed, the *hrpB* mutant strain was found to be non-pathogenic and the *phcA* mutant strain was observed to be highly reduced for virulence. The *pilT* mutant was reduced for virulence but more virulent than the *phcA* mutant. In case of *rpoN2* mutant strain, the virulence proficiency was found to be similar to the wild type *R. solanacearum* F1C1 (Figure 8). The virulence phenotypes and disease progression of *hrpB, phcA, pilT* and *rpoN2* mutants were in concordance with the virulence phenotype data of tomato seedling (mentioned in previous chapter) as well as in grownup plant as reported in literature (Ray et al., 2015).

**Figure 8:**
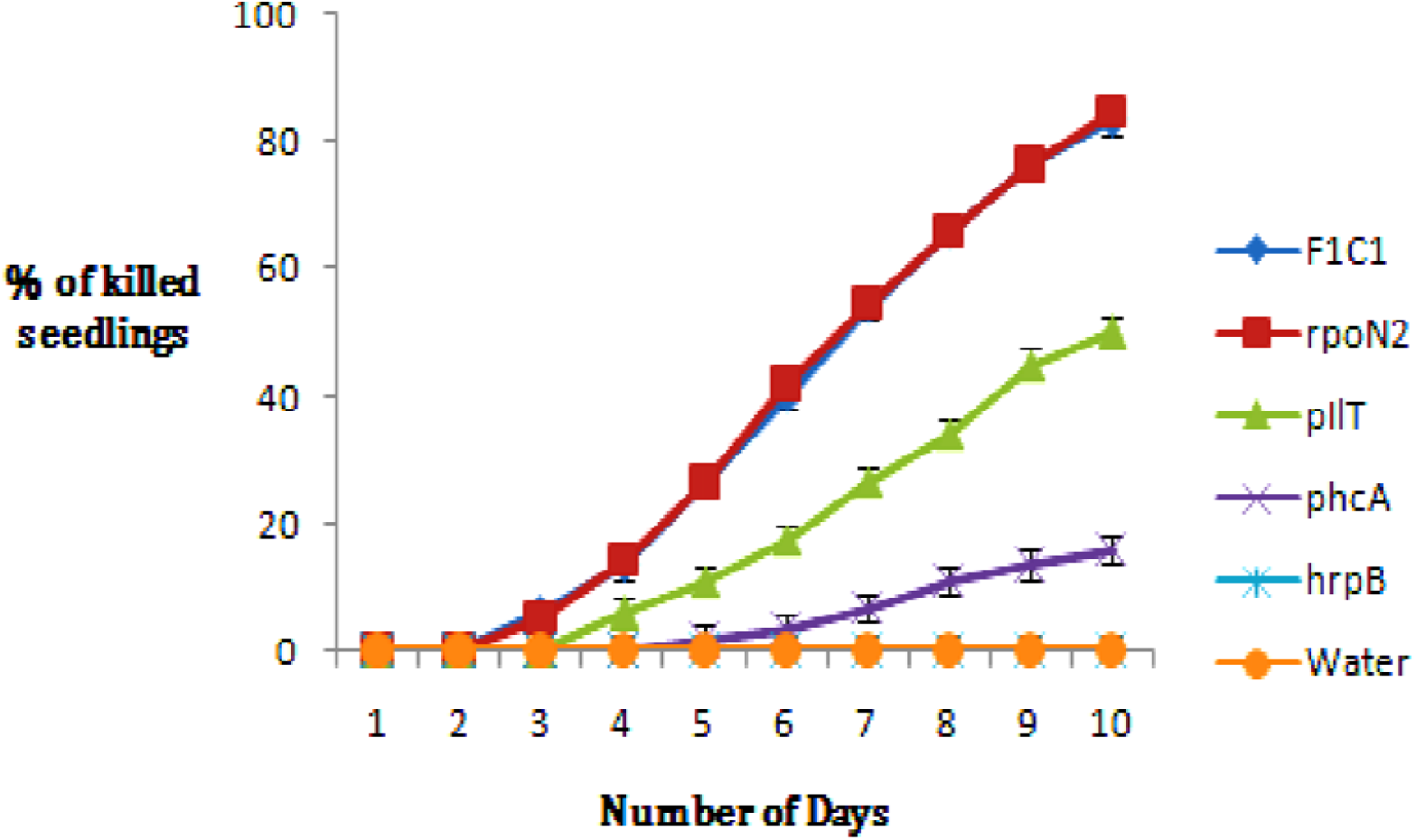
Comparative pathogenicity assay of F1C1 and other derivative mutant strains in eggplant seedlings. This graph is showing the pathogenicity comparison of F1C1 to its derivative mutants*rpoN2, hrpB, phcA* and *pilT*. The number (%) of killed seedlings in case of *phcA* and *pilT* were much lesser than the wild type (F1C1). While in case of *hrpB*, no wilting was observed similar to water control set. But in case of *rpoN2*, there was no affect of pathogenicity similar to wild type F1C1.

### 4.4 Wrapping of thin layer cotton fiber around eggplants seedling root to study its impact on *R. solanacearum* pathogenicity

During the standardization of pathogenicity assay in eggplant seedlings, it was observed that if a thin layer of cotton fiber was wrapped around the root of eggplant seedlings just after the pathogen inoculation, wilting and disease progression was found to be enhanced in comparison to infected eggplant seedling (sets) without cotton wrapping (Figure 9 & 10).

**Figure 9:**
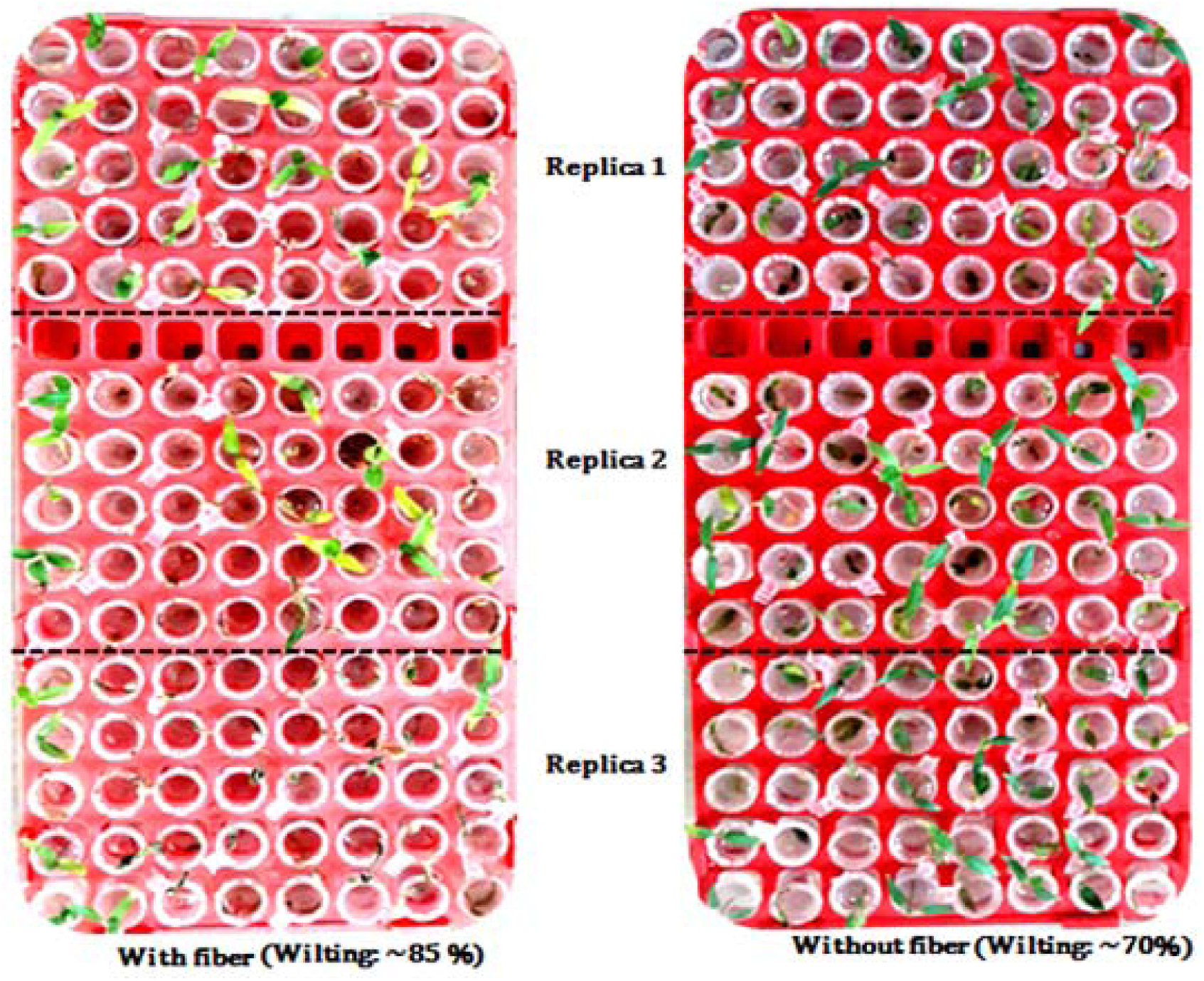
Comparative pathogenicity assay between wrapped and unwrapped eggplant seedlings. The picture in the left side(cotton wrapped)with a set of seedling are exhibiting more number of wilting in comparison to set of infected seedlings in the right side (unwrapped) picture. In addition to that, seedling leaves in the left sidepicture are yellowish in comparison to rightside which is also supporting pathogenicity enhancement in cotton wrapped seedlings.

**Figure 10:**
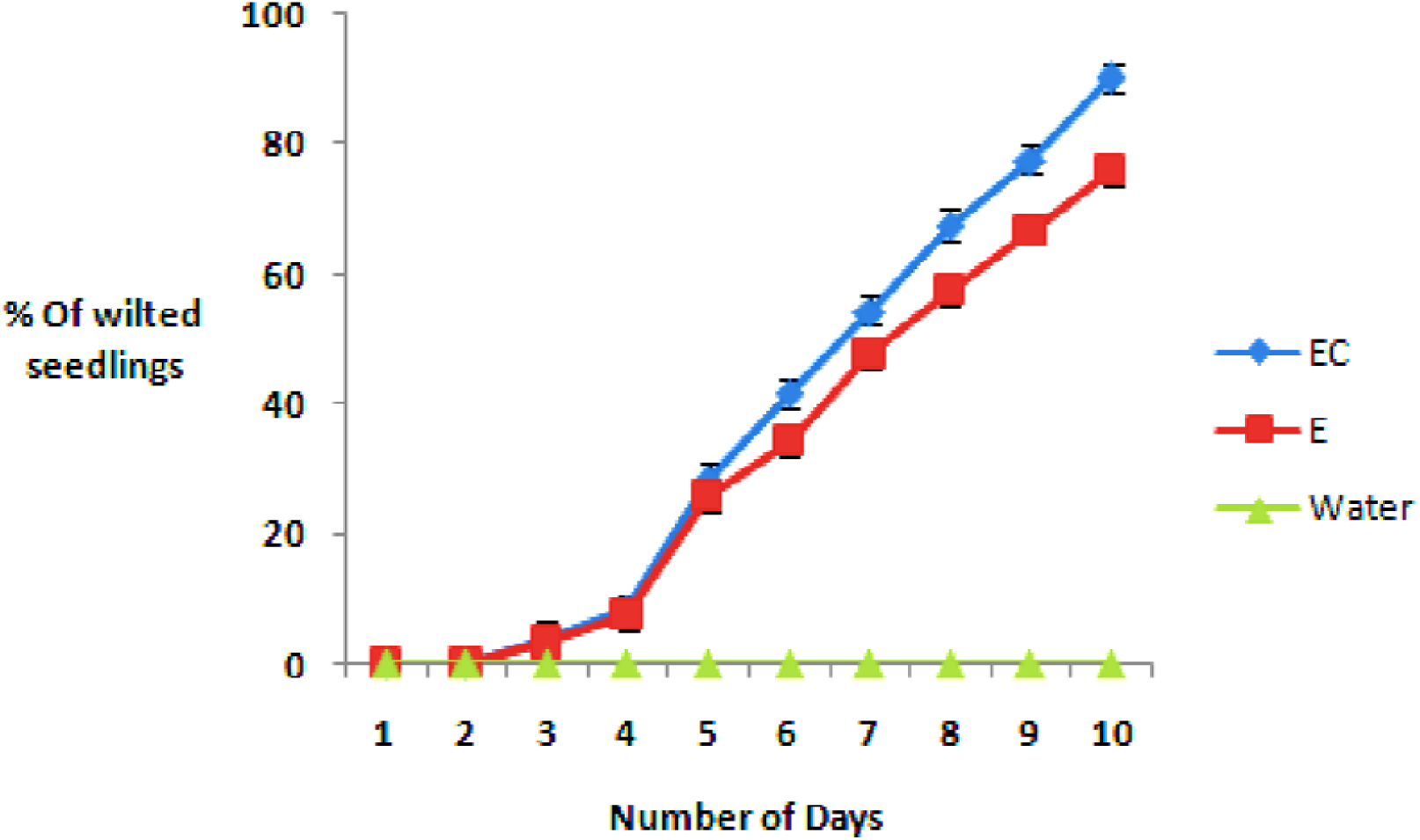
Pathogenicity assay in cotton wrapped and unwrapped eggplant seedlings(statistical analysis : Appendix II) This picture is showing the pathogenicity assay of F1C1 in 2 (two) different sets of an eggplant cultivar by root inoculation method. In this graph, EC is indicating eggplant with the wrapping cotton fiber while E is indicating the eggplant without the wrapping of cotton fiber. This graph is clearly showing that through this method of inoculation, F1C1 is significantly more virulent and disease progression (5^th^ day onwards) in case of EC or cotton fiber wrapped eggplant seedlings.

The effect of cotton wrapping around the root of eggplant seedling was an interesting observation of this study. But to understand and reveal the exact reason behind the enhancement of virulence and pathogenicity, there is a necessary of further dipper investigation of plant-microbe interaction behavior during early stage of infection and attachment.

## 5 Discussion

In this work, we are documenting the standardization of a protocol to study *R. solanacearum* pathogenicity (by root inoculation approach) in 10-12 days old eggplant seedlings under gnotobiotic and additional nutrients free conditions. This standardized protocol was found to be very efficient and consistent for the pathogenicity assay at seedlings stage of eggplant. In addition to that, it has also shown the reproducibility in other different eggplants cultivars. This newly standardized protocol is successfully validated with the help of known mutant gene function of *R. solanacearum* derivatives such as *hrpB, phcA* and *pilT* mutant strains. Over and above, this inoculation process is also found to be easy to investigate the pathogen colonization study at seedling stage. Colonization study of this bacterial pathogen after post inoculation clearly manifested that the pathogen is successfully associated with the root and shoot regions of the infected seedlings.

However, this pathogenicity standardization protocol in eggplant seedlings is superficially looks similar to tomato seedling, but actually it has many minute differences. In case of eggplants seedlings, pathogenicity symptom appearance, seed germination process and seedling age were very different from tomato seedlings. It took lot of time and effort to optimize optimum germination of seed as well as disease progression and wilting in eggplant seedlings. In this protocol, seed soaking time in the presence of distilled water for eggplant seeds was optimized as 5-6 days while in case of tomato, 2 days of soaking duration was found to be sufficient to get optimum germination. The exact reason behind the requirement of longer soaking duration for optimum germination about eggplant seeds is yet to know. During standardization, we felt that eggplant seed coat was harder and thicker than that of tomato, which might be a probable reason behind requirement of longer soaking duration of eggplant seeds for optimum germination. The other point was that the initial disease symptom appearance in case of eggplant seedling was little bit late in compression to tomato seedling and the assay got completed by 10 days in eggplant compared to 7-8 days in case of tomato seedling. Over and above it is also important to mention seedling age of eggplant at the time of pathogen inoculation, which was different than tomato: 10-12 days old eggplant seedlings were used for pathogen inoculation and disease progression study as compared to 6-7 days old seedlings of tomato. Therefore, because of several differences of this standardized protocol from the earlier reported protocol for tomato seedlings, we are documenting this study as a separate protocol for pathogenicity assay in eggplant seedling. This standardized protocol for pathogenicity assay is also found to be very much consistent and reproducible in other three different eggplant cultivars. Hence, on the basis of reproducibility and consistency of this protocol, this study is suggesting the carried out protocol may be an efficient way to study plant-microbe and wilt disease progression of *R. solanacearum* at the early seedling stage of eggplant. In addition to that, the pathogenicity assay can be performed by maintaining near to natural mode of inoculation in less resource dependent manner. It is pertinent to note that this is the first standardization report on *R. solanacearum* pathogenicity study in cotyledon stage seedlings of eggplant by root inoculation.

Very interestingly, in-between hit and trial of pathogenicity assay standardization, this study have noticed that wrapping of a thin layer of sterile cotton fiber around the root of the pathogen inoculated eggplant seedling used to enhance the wilting and disease progression of *R. solanacearum*. In an initial study we also observed that poly ethylene fiber was enhancing the *R. solanacearum* pathogenicity in eggplant seedlings like that of the cotton fiber. This observation might be important to know the impact fiber structures present in fields for the natural infection *R. solanacearum* in eggplant cultivars and other hosts. Although exact mechanism is still unknown, probably this cotton fiber used induced some adhesion functions such as type IV pili during initial attachment and/or induced biofilm formation pathway to stimulate additional virulence and wilting in the eggplant seedling (Liu et al., 2001; Siri et al., 2014; Persat et al., 2015).We have not come across any references regarding the impact of fibers on *R. solanacearum* pathogenicity in host plants. In future, it will be interesting to investigate and understand the exact mechanism and correlation of cotton fiber in disease progression and wilting.

## 6. Acknowledgment

Niraj Singh is thankful to Tezpur University, Assam, India as well as Royal Global University, Assam, India for providing an opportunity to work. Over and above, Niraj Singh is also very much thankful to Prof. Suvendra Kumar Ray, Tezpur University, for motivation and kind support. In addition to that, Niraj Singh also thankful to Department of Biotechnology, Government of India for the DBT JRF and SRF fellowship and providing necessary facilities.

